# Amino acid auxotrophies in human gut bacteria are linked to higher microbiome diversity and long-term stability

**DOI:** 10.1101/2023.03.23.532984

**Authors:** Svenja Starke, Danielle MM Harris, Johannes Zimmermann, Sven Schuchardt, Mhmd Oumari, Derk Frank, Corinna Bang, Philip Rosenstiel, Stefan Schreiber, Norbert Frey, Andre Franke, Konrad Aden, Silvio Waschina

## Abstract

Amino acid auxotrophies are prevalent among bacteria. They can govern ecological dynamics in microbial communities and indicate metabolic cross-feeding interactions among coexisting genotypes. Despite the ecological importance of auxotrophies, their distribution and impact on the diversity and function of the human gut microbiome remain poorly understood. This study performed the first systematic analysis of the distribution of amino acid auxotrophies in the human gut microbiome using a combined metabolomic, metagenomic, and metabolic modeling approach. Results showed that amino acid auxotrophies are ubiquitous in the colon microbiome, with tryptophan auxotrophy being the most common. Auxotrophy frequencies were higher for those amino acids that are also essential to the human host. Moreover, a higher overall abundance of auxotrophies was associated with greater microbiome diversity and stability, and the distribution of auxotrophs was found to be related to the human host’s metabolome, including trimethylamine oxide, small aromatic acids, and secondary bile acids. Thus, our results suggest that amino acid auxotrophies are important factors contributing to microbiome ecology and host-microbiome metabolic interactions.

## Background

The metabolic processes performed by the human gut microbiota have a crucial impact on human metabolism and health(1–3). For instance, various human gut bacteria produce the short-chain fatty acid butyrate. Butyrate is a primary energy source for human colonocytes(1) and intersects with host immunological processes by mediating anti-inflammatory effects(4,5). Another notable metabolic interaction between the human host and its gastrointestinal microbiota is the microbial transformation of aromatic amino acids into various metabolites. Recent studies suggest that aromatic amino acid-derived metabolites such as the auxins indole-3-propionic acid and indole-3-acetic acid can modulate the host immune system(6,7). Thus, these and several further studies provide evidence that gut microbial metabolites are essential factors in the pathophysiology of inflammatory diseases and the efficacy of immunomodulatory therapies(7–10).

The repertoire of molecules synthesized and eventually released by individual gut microbes comprises metabolic by-products that serve the dual purpose of energy metabolism and facilitating the biosynthesis of essential metabolites necessary for cellular maintenance and proliferation. However, often not all metabolites required for growth and survival (i.e., nucleotides, vitamins, amino acids) can be de-novo synthesized by gut-dwelling microorganisms, rendering those organisms dependent (termed *auxotrophic*) on the uptake of the focal metabolite from the microbial cell’s nutritional environment. Several *in silico* studies have applied genome-mining approaches, suggesting that most analyzed gut bacteria lack biosynthetic pathways for producing at least one proteinogenic amino acid(11,12) or a growth-essential vitamin(13,14). In addition, *in vitro* growth experiments have confirmed specific amino acid and vitamin auxotrophies in common human gut bacteria(13,15,16).

The prevalence of auxotrophs in the human gut microbiome raises the question of the source of the required metabolites in the gastrointestinal growth environment. There are three potential sources of essential nutrients for microbial growth: (i) Required metabolites could be diet-derived. However, amino acids and vitamins are usually efficiently absorbed by the human host in the small intestine(17), limiting the accessibility of diet-derived essential nutrients for the majority of the gut microbial community, which resides in the colonic region(18). (ii) Metabolites required by auxotrophic microorganisms in the gastrointestinal tract may be host-derived, e.g., from proteins and peptides secreted by the gut epithelium into the gut lumen or from apical proteins of the host epithelial cell layer accessible to gut microorganisms(15). (iii) Auxotrophic members of the gut microbial community might obtain essential nutrients via cross-feeding interactions with prototrophic organisms within their microbial community(19,20).

While the exchange of electron donor metabolites (e.g., acetate-or lactate cross-feeding) between different microorganisms is well-documented for the human gut microbiome(21–23), the extent of cross-feeding interactions via the exchange of essential nutrients such as amino acids and vitamins remains still unknown. However, in vitro experiments of synthetic microbial communities suggest that co-cultured microorganisms, which are auxotrophic for different compounds, can support each other’s growth by exchanging the focal metabolites(24). Furthermore, theoretical ecological models suggest that cross-feeding interactions between auxotrophic organisms within complex communities can increase community diversity through metabolic niche expansion(25) and community robustness to ecological perturbance(26), such as changes in the composition of the chemical environment. Thus, cross-feeding of amino acids and vitamins between different members of the human gut microbiota could be crucial determinants of microbiome dynamics, resilience, and the contribution of gut microbes to human metabolism and health.

In this study, we applied genome-scale metabolic modeling to predict the distribution and diversity of amino acid auxotrophies in the human gut microbiome. The predictions were combined with stool metagenomic sequencing and targeted serum metabolomics from observational human cohort studies to estimate auxotrophy frequencies and their impact on the human metabolome. We found that amino acids that are essential to the human host are also the most common auxotrophies in the human gut microbiome. Intriguingly, a higher frequency of auxotrophies was associated with long-term stability of the microbiome community composition. Furthermore, a higher number of auxotrophies among gut bacteria was associated with higher diversity of the gut bacteria and increased levels of aromatic compounds of putative microbial origin in the human serum metabolome.

## Results

### Prediction and validation of auxotrophies with genome-scale metabolic modeling

To estimate the overall distribution of amino acid auxotrophies in the human gut microbiome, we predicted the amino acid production capacities using genome-metabolic modeling for all bacterial genomes (n=5 414) from the ‘Human Reference Gut Microbiome (HRGM)’ collection(27). Auxotrophies were predicted for the 20 proteinogenic amino acids by comparing the model’s growth with and without the amino acid using flux-balance analysis. If the model was not able to grow without the amino acid, then an auxotrophy was predicted (Fig. 1). To exclude an overprediction of auxotrophies due to genome incompleteness, we correlated the genome completeness and the number of auxotrophies predicted. Results showed a negative relationship between genome completeness and the number of auxotrophies per genome (Supplementary Fig. S1, ρ=-0.50, p≤2.2e-16). To combat this, the genomes were filtered for completeness ≥85% and contamination ≤2%. Only the filtered metabolic models (n=3 687) were used to predict auxotrophies and ongoing analysis. All auxotrophies predicted for HRGM models are in the supplementary material (Supplementary Table S1).

**Figure 1.**
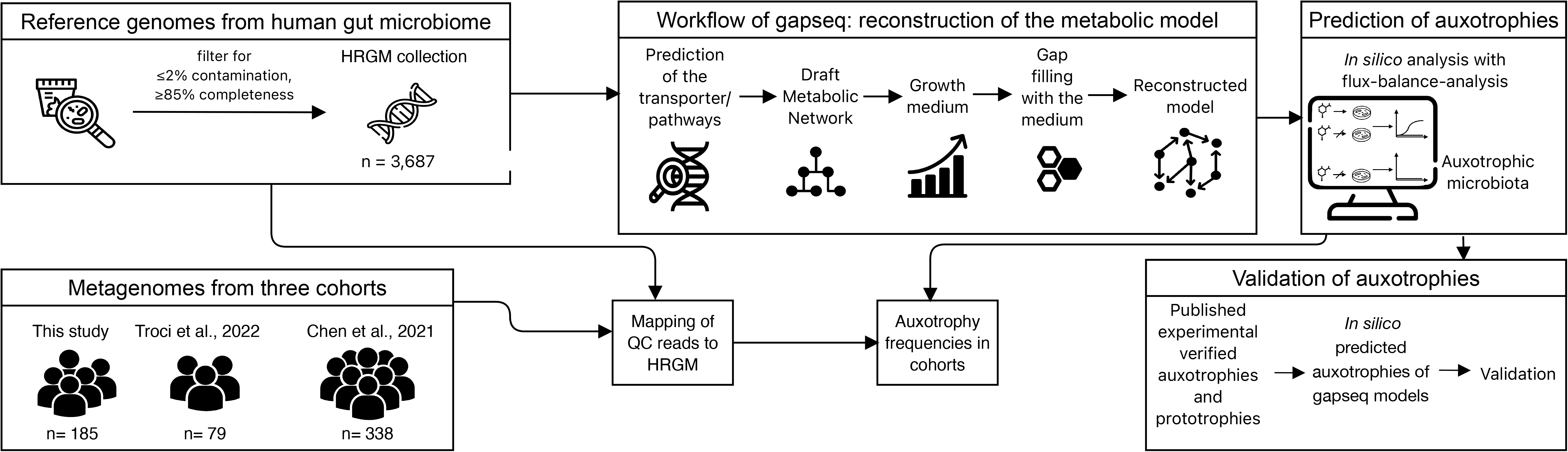
Workflow for the prediction of auxotrophies with genome-scale metabolic modeling. Gapseq was used to reconstruct genome-scale metabolic models from genomes of the Human Reference Gut Microbiome (HRGM) catalog(27). The workflow of gapseq to reconstruct metabolic models consists of five steps: transporter/metabolic pathway prediction, draft metabolic network construction, growth medium prediction, gap filling, final model reconstruction. Auxotrophy prediction was performed using flux-balance analysis and validated by reconstructing gapseq models from experimentally verified auxotrophic strains. The predicted auxotrophies were compared on strain level from gapseq and AGORA2 models to experimentally verified auxotrophies. QC reads of cohorts were mapped on HRGM. Auxotrophy frequencies in cohorts were determined by mapping QC reads from the metagenomes of the cohorts to genomes from HRGM collection. Icons are from www.flaticon.com (creators: photo3idea_studio, Freepik, surang, Eucalyp, Voysla, juicy_fish, smashingstocks, SBTS2018, creative_designer).

A recent study has reported discrepancies between *in silico* predictions using metabolic models reconstructed with carveme (28) and in vitro studies of amino acid auxotrophies in bacteria(13). To validate our gapseq-based auxotrophy predictions, we compared the predictions on strain level with *in vitro* experimentally verified auxotrophies as reported in previous studies for a total of 36 gut bacteria (Supplementary Table S2), of which most were already summarized by Ashniev et al. 2022 (29). If a genome assembly of the experimentally tested strain was available on NCBI RefSeq, we reconstructed the genome-scale metabolic model and predicted the auxotrophies. In addition to auxotrophy predictions using our gapseq model collection (Supplementary Table S1), we also tested models from the AGORA2 collection (Supplementary Table S3). Auxotrophy predictions using gapseq models had a sensitivity of 75.5%, a specificity of 95.9%, and an accuracy of 93%. The auxotrophies predicted by the AGORA2 models showed a lower degree of agreement with the experimental data: sensitivity (43.4%), specificity (92.3%), and accuracy (81.7%). In addition, we reconstructed genome-scale metabolic models for 124 bacterial genotypes known to be prototrophic for all 20 proteinogenic amino acids(28) to further validate our auxotrophy predictions (Supplementary Table S4). We note that the 124 prototrophic genotypes are isolates from diverse isolation sources and not from the human gut. However, the resource can be used to estimate the rate of false auxotrophy predictions(28). In total, 99.1% of all predictions coincided with the known amino acid prototrophies of the organisms, thus suggesting a false auxotrophy prediction rate of less than 1%. In general, the frequency of auxotrophy predictions among genomes from human gut bacteria is generally higher compared to the collection of 124 prototrophic genomes (Supplementary Fig. S2), indicating that the high frequency of auxotrophies cannot be explained by a false-positive rate associated with potential pitfalls in the model reconstruction workflow.

### Amino acid auxotrophies are common in the human gut microbiome

Auxotrophies for tryptophan were the most prevalent, at 63.9% of the genomes in the HRGM catalog(Fig. 2). Isoleucine, leucine, and valine (BCAA, branched-chain amino acids) auxotrophies were also detected with a high abundance (40.1%, 40%, 41.1%, respectively). No auxotrophies were detected for alanine, aspartate, and glutamate. We further analyzed the observed auxotrophies at the taxonomy level by comparing the proportion and number of auxotrophies on phylum and order level (Supplementary Fig. S3). Actinobacteriota were shown to have a higher proportion of BCAA auxotrophies compared to prototrophies (Supplementary Fig. S3). For tryptophan, a higher proportion of auxotrophic to prototrophic bacteria was observed in Firmicutes, Actinobacteriota, and Fusobacteriota. Fusobacteriota generally had a higher auxotrophic to prototrophic ratio for almost all amino acids, whereas the opposite was predicted for Proteobacteria. This observation is further supported by the number of auxotrophies found per genome for Proteobacteria and Fusobacteriota (Supplementary Fig. S4). Additionally, the results suggest that auxotrophic genotypes have lost the genes for most of the enzymes required for the biosynthesis of the focal amino acid (Supplementary Fig. S5).

**Figure 2.**
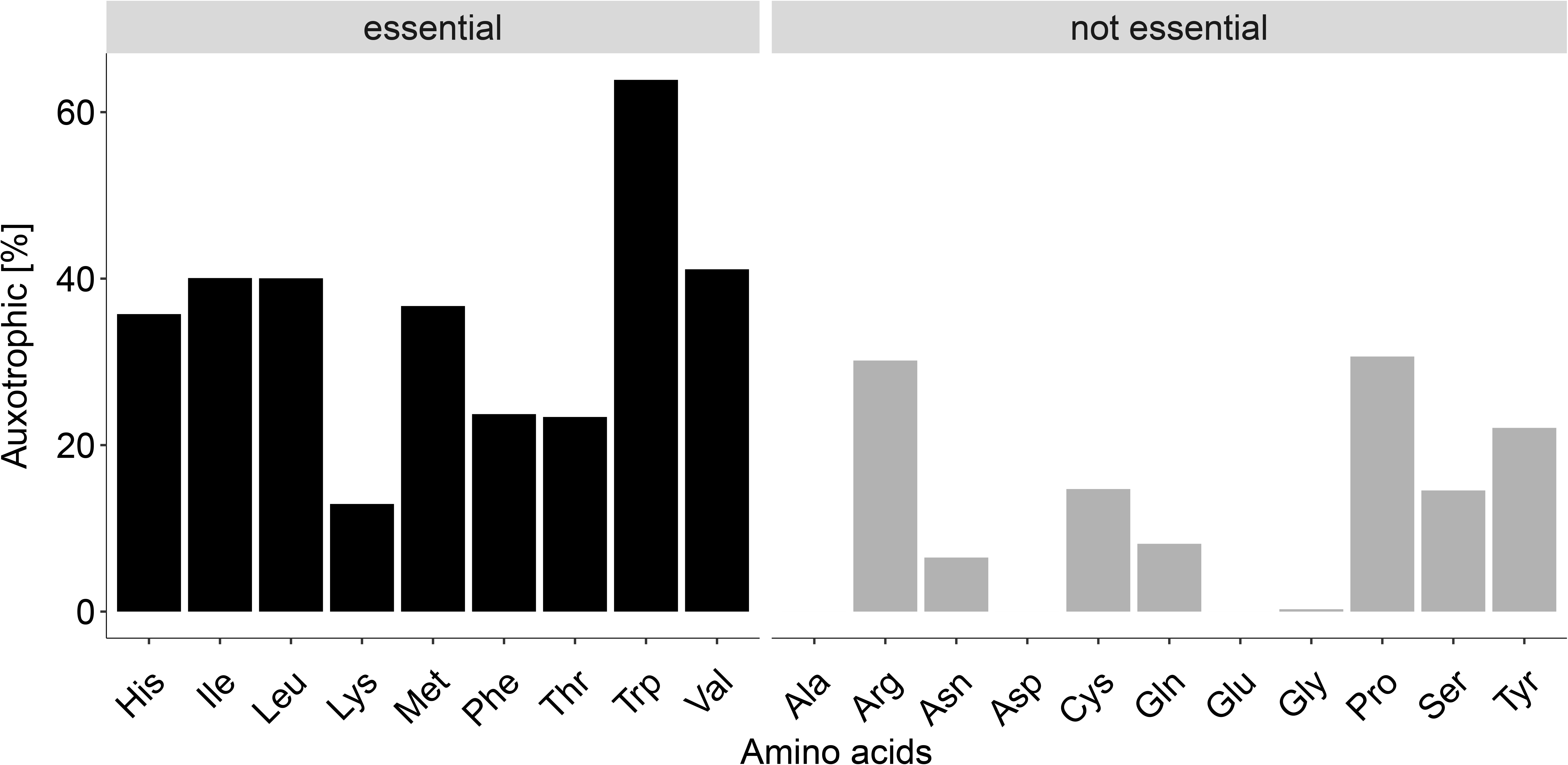
Abundances of auxotrophies in 3 687 genomes. The predicted amino acid auxotrophies in HRGM genomes were categorized into human essential and non-essential amino acids.

Taken together, the results indicate that amino acid auxotrophies are prevalent in the human gut microbiome.

### Amino acid auxotrophies are associated with the profile of fermentation products

Amino acid biosynthesis pathways and pathways producing fermentation products share common precursor metabolites (Fig. 3B). For example, pyruvate is a central metabolite that is utilized for the biosynthesis of the BCAA as well as in some gut bacterial species for lactate formation, underlining the interconnection of amino acids and energy metabolism in the metabolic network.

**Figure 3.**
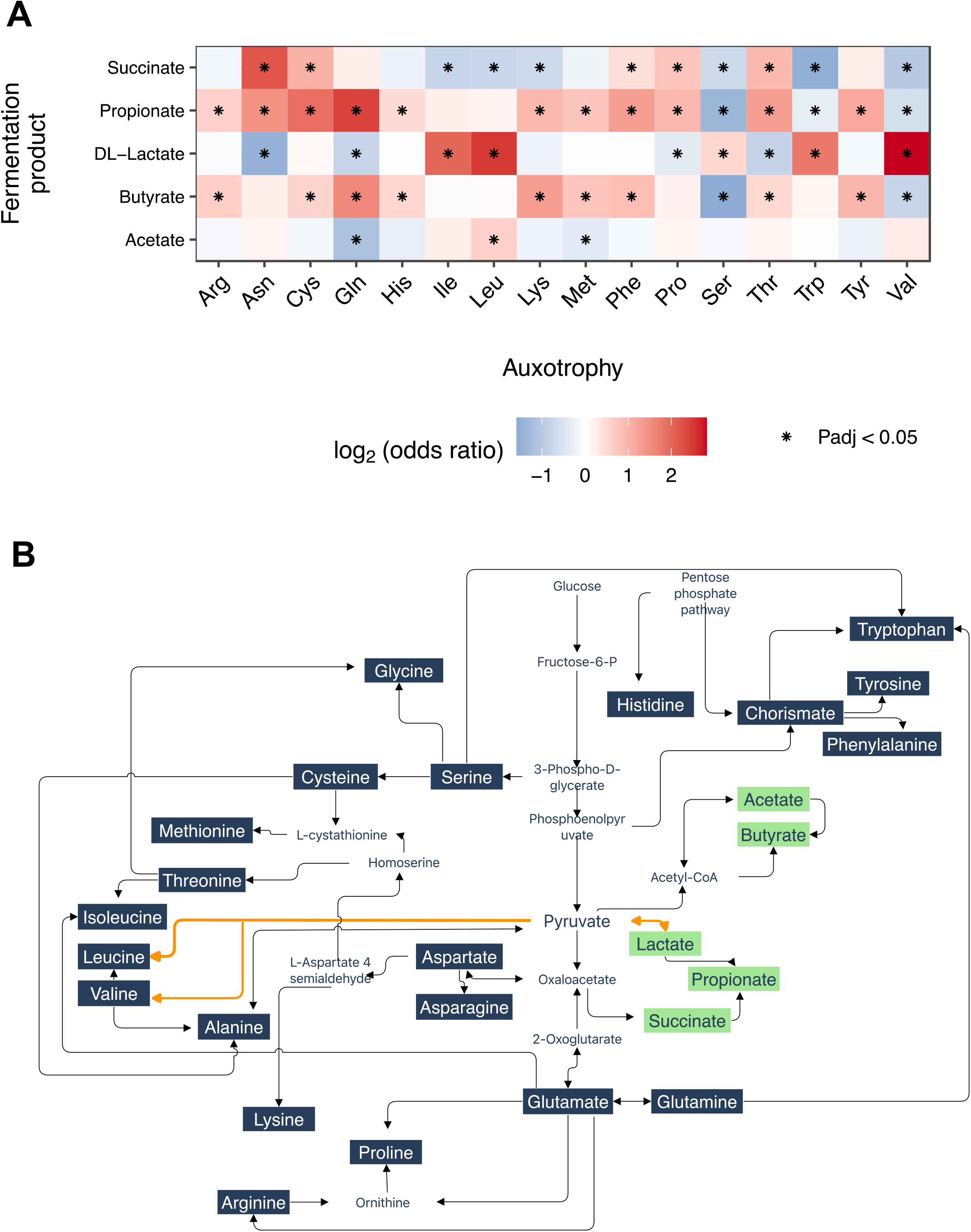
Associations of auxotrophies and fermentation products. (A) Comparison of fermentation product production rates in auxotrophic and prototrophic bacteria. Production rates of fermentation by-products were predicted with flux-balance analysis (cutoff-value > 1 mmol/gDW) in 3 687 HRGM genomes. The association with the auxotrophic or prototrophic phenotype was statistically evaluated with the Fisher test for exact count data by calculating odds ratios. Asterisks denote FDR-corrected p-values <0.05. (B) Interconnection between the pathways of formation of fermentation products and amino acids, based on MetaCyc pathways(76). We note that not all the displayed reactions/pathways occur in every gut bacterial genotype. The metabolic network shown displays pathways commonly found in human gut metagenomes and linked to amino acid biosynthetic pathways.

Here, we investigated whether bacteria that are auxotrophic for specific amino acids are commonly associated with specific profiles of fermentation products. Therefore, we predicted the metabolic by-products of cell growth and compared those results with the auxotrophy predictions for the corresponding organisms (Fig. 3A). BCAA auxotrophic bacteria were more likely to produce lactate in comparison to prototrophic bacteria (Fisher’s exact test for count data,-log_2_(Odds Ratio (OR)) = 2.0-2.8, FDR-corrected p-value<0.05). Propionate production was commonly predicted for glutamine auxotrophic gut bacteria (-log_2_(OR)= 2.4, FDR-corrected p-value<0.05) and by cysteine auxotrophs (-log_2_(OR)= 1.9, FDR-corrected p-value <0.05). Succinate is predominantly produced by asparagine auxotrophic gut bacteria (-log_2_(OR)= 2.2, FDR corrected p-value <0.05). For butyrate, there was a higher association with glutamine auxotrophic bacteria (-log_2_(OR)= 1.6, FDR-corrected p-value <0.05).

The association of auxotrophic bacteria with the production of organic acids might be explained by the distribution of reaction fluxes through the metabolic network. For instance, pyruvate is a metabolic precursor for the *de novo* biosynthesis pathways for BCAA but also for lactate formation (Fig. 3B). Pyruvate not used for BCAA biosynthesis in auxotrophic genotypes might be redirected towards lactate production. Thus, our findings suggest a plausible interplay in resource allocation between a microorganism’s energy metabolism strategy and its auxotrophy profile.

### More diverse gut microbiomes are characterized by a higher auxotrophy frequency

To estimate the frequency of auxotrophies in the gut microbiome of individual persons, we quantified the relative abundance of gut bacterial genotypes from the HRGM catalog using stool metagenomes of 185 healthy adults. As mentioned above, we found a negative correlation between the number of auxotrophies and genome completeness levels (Supplementary Fig. S1). To validate that higher genome completeness levels do not affect the general pattern in the auxotrophy distribution of individual microbiomes, we determined auxotrophy frequencies with different cutoff values for completeness (80%-95%) of the reference genomes used for quantification. Overall, the distribution of auxotrophy frequencies remained robust to increasing genome completeness levels (Supplementary Fig. S6). Therefore, we decided to keep the 85% completeness level described above.

Strikingly, auxotrophies for amino acids that are essential to the human organism were more frequent than non-essential amino acids (Fig. 4A). The highest percentage of bacteria were auxotrophic for tryptophan, followed by isoleucine and histidine (median: 54%, 28.7%, 28%, respectively). Auxotrophies for leucine, methionine, phenylalanine, arginine, and valine were found with a median frequency of >20% (Fig. 4A). The lowest frequencies were detected for serine, lysine, asparagine, aspartate, alanine, and glutamate auxotrophies. Additionally, we were interested in the relationship between the proportion of auxotrophic bacteria in the human gut and the overall microbiome diversity calculated as the Shannon index (Fig. 4B-C). Overall, increasing frequencies of almost all amino acid auxotrophies are accompanied by increasing microbiome diversity (Spearman correlation, Fig. 4B). Further, we correlated the Shannon diversity with the abundance-weighted average of the number of auxotrophies per metagenome sample, which takes the relative abundance of each genome and its total number of amino acid auxotrophies into account. With an increasing number of auxotrophies, an increase in the diversity was observed (Fig. 4C, ρ=0.27, p=0.00018). This result may point towards a positive influence of auxotrophic bacteria on the microbial diversity in the gut, presumably via a higher degree of amino acid cross-feeding interactions between genotypes that are auxotrophic for different amino acids. To test this, we calculated the pairwise dissimilarity (Hamming distance) between the binary auxotrophy profiles of genomes and the means of those differences per metagenome sample as an indicator for potential cross-feeding in the respective gut microbial community. An increasing average Hamming distance was positively associated with increased gut diversity (Fig. 4D, ρ=0.32, p=0.00001). Overall, a higher number of auxotrophies in the gut community is positively correlated with a higher diversity.

**Figure 4.**
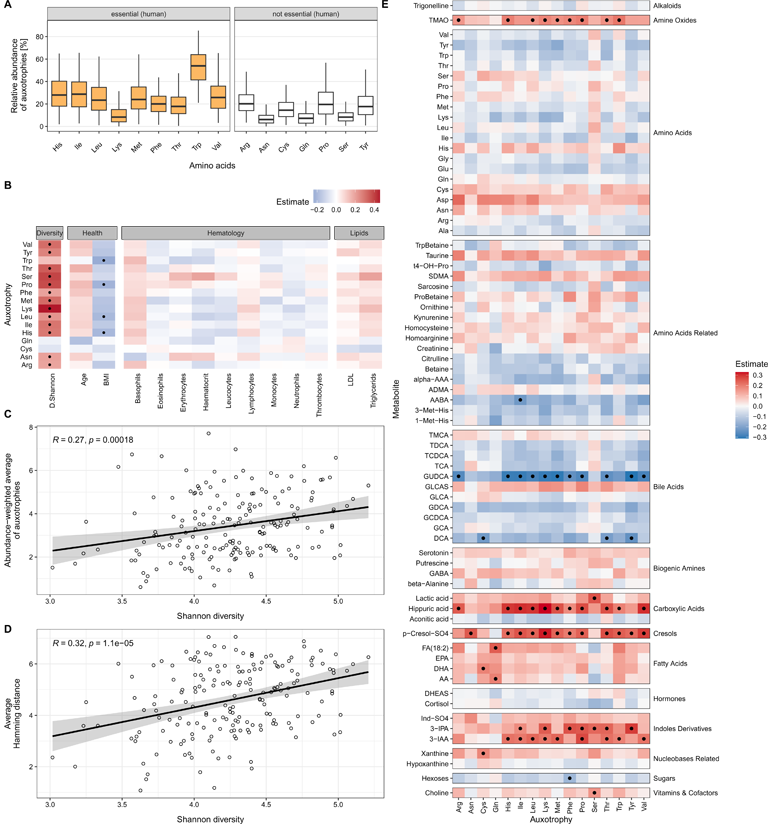
Distribution of auxotrophies in human gut microbiomes from 185 healthy adults, their association with diversity, and serum metabolite levels. (A) Boxplots display the abundance of amino acid auxotrophies in the human gut microbiome (n=185 samples). (B) Partial Spearman correlation between the frequency of auxotrophic gut bacteria and serum levels of health markers and microbiome Shannon diversity. Dots indicate significant associations (FDR-corrected p-values < 0.05, adjusted for the potential confounders age, sex, and BMI). (C) The abundance-weighted average of auxotrophies was calculated and correlated with the Shannon diversity (Spearman correlation, ρ =0.60, p<2.2e-16). (D) The average hamming distance was calculated to study the metabolic dissimilarity of auxotrophy profiles of coexisting genotypes and, therefore, potential cross-feeding interactions within the microbial communities. With the Spearman correlation, the association between the calculated average hamming distance and the Shannon diversity in the gut was estimated (ρ =0.62, p<2.2e-16). (E) Partial Spearman correlations between the serum levels of metabolites and the frequency of auxotrophic bacteria in the gut microbiome. Abbreviations for the serum metabolite levels can be found in Supplementary Table S5. Dots indicate significant associations (FDR-corrected p-values < 0.05, adjusted for confounders age, sex, and BMI).

### Associations of gut bacterial auxotrophies for amino acids with host health markers and the serum metabolome

The involvement of microbial metabolism in host health has been examined in several other studies (30,31) but not yet for the frequency of gut microbial amino acid auxotrophies. Our results showed that several amino acid auxotrophic bacteria are inversely associated with the stool donor’s BMI (Fig. 4B, partial Spearman correlation). No statistically significant associations with blood cell counts were found (Fig. 4B). Additionally, we correlated targeted metabolomics data from serum samples with the frequencies of specific amino acid auxotrophies (Fig. 4E, partial Spearman correlation). Positive correlations were found between the tryptophan-derived 3-indoleacetic acid (3-IAA) as well as 3-indolepropionic acid (3-IPA) and tryptophan auxotrophic gut bacteria. Additionally, several other amino acid auxotrophies showed positive correlations with these metabolites. P-cresol sulfate was positively correlated with many amino acid auxotrophies. Further, several significant associations were detected with metabolites from bile acid metabolism. Negative correlations were observed for glycoursodeoxycholic acid (GUDCA), a conjugated secondary bile acid metabolite, and several amino acid auxotrophies. Further, negative correlations with the bile acid metabolite deoxycholic acid (DCA) were found for the frequencies of tyrosine, threonine, and cysteine auxotrophies. Positive associations were also observed for hippuric acid and TMAO with several amino acid auxotrophies. Interestingly, no significant associations were found for serum levels of amino acids and amino acid-related compounds (Fig. 4E).

Taken together, the frequency of auxotrophic bacteria is related to serum levels of several metabolites. The gut microbial contribution to serum metabolite levels was predominantly found for metabolites previously reported to be of microbial origin (e.g., 3-IAA) or derived from gut microbially-produced compounds (e.g., TMAO).

### Analysis of longitudinal microbial composition data suggests a positive influence of auxotrophies on gut microbiome stability

So far, our results suggest an involvement of auxotrophic bacteria on the gut microbial diversity. Based on this observation, we further wanted to analyze whether the frequency of auxotrophies also impacts the microbiome’s long-term stability using data from two longitudinal studies. Therefore, we re-analyzed recently published metagenomic data from two human cohort studies (32,33). Troci et al. included two stool metagenomes from 79 healthy individuals each where stool samples were three years apart(32). The longitudinal study of Chen et al. involved two stool metagenomes from 338 individuals with a time difference between samples of four years(33). Microbiome stability over the time periods was assessed by calculating the UniFrac distance for the microbial composition between the two time points for each participant. Since the UniFrac distance ranges between 0 (lowest possible dissimilarity) and 1 (highest dissimilarity), we calculated the inverse values (1-UniFrac) as a microbiome stability measure. The abundance-weighted average of auxotrophies per genotype was positively correlated with microbiome stability in both cohorts (Fig. 5A, Spearman rank sum correlation test, Troci et al.: ρ=0.31, p=0.006, n =79; Chen et al.: ρ=0.14, p=0.0094, n = 338). We also correlated individual amino acid auxotrophy frequencies with microbiome stability to understand the impact of individual amino acid auxotrophies on long-term stability. A statistically significant positive correlation was found in both cohorts for many amino acid auxotrophies, while no negative correlation was observed (Fig. 5C).

**Figure 5.**
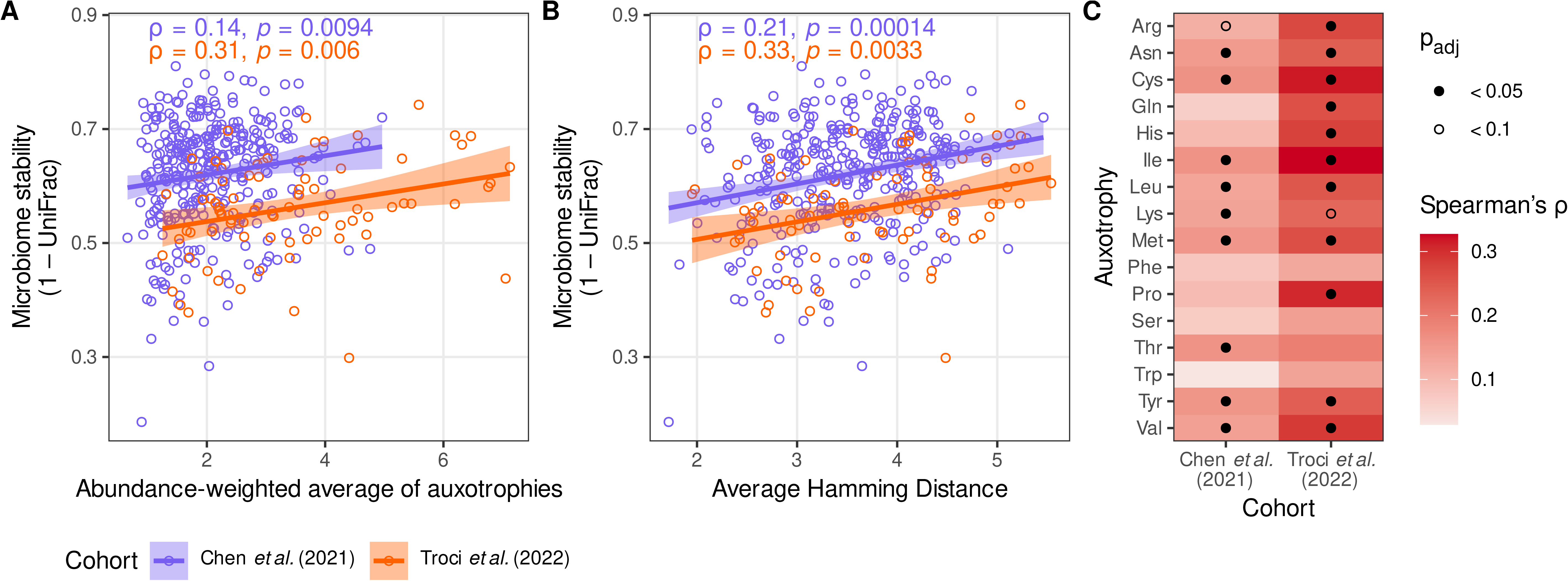
Influence of auxotrophies on long-term stability of the human gut microbiome. (A) The stability of the human gut microbiome was calculated as 1 minus the UniFrac distance between the two time points in the longitudinal studies and correlated with the abundance-weighted average of auxotrophies at the first time point to study a potential influence of auxotrophies on the long-term stability of the human gut microbiome. (B) The average Hamming distance was calculated for the first time point and then correlated with the 1-UniFrac value to investigate the influence of potential cross-feeding on long-term stability. (C) The contribution of individual amino acid auxotrophies on the stability was calculated with the Spearman correlation between the 1-UniFrac values and individual amino acid auxotrophy frequencies.

Next, long-term microbiome stability was also tested for a statistical association with the average Hamming distance with samples, which represents a measure of the dissimilarity between the auxotrophy profile of co-existing genotypes and a potential indicator for the degree of amino acid cross-feeding in the microbial community. A notable positive correlation was observed for the average Hamming distance with microbiome stability in both cohorts (Fig. 5B, Troci et al.: ρ=0.33, p=0.0033, n =79; Chen et al.: ρ=0.21, p=0.00014, n = 338.), suggesting a potential positive impact of amino acid cross-feeding among auxotrophy genotypes on the long-term stability of microbiome composition. Auxotrophic bacteria have a high dependence on their nutritional environment. Here, we wanted to test if a higher dietary intake of amino acids affects the relative abundance of amino acid auxotrophic bacteria in the gut. Therefore, we used the dietary intake data obtained from food frequency questionnaires from Troci et al.(32). For both study time points, the intake of amino acids was tested for correlation with the frequency of amino acid auxotrophies in the microbiomes. No significant correlations between the frequency of auxotrophic bacteria and the dietary intake of amino acids were observed (Supplementary Fig. S7).

In sum, our results suggest a positive effect of auxotrophies on gut microbiome stability. Further, the data suggest that amino acid cross-feeding may contribute to the compositional stability of the gut microbiome. Surprisingly, we found no evidence of diet’s effect on auxotrophy frequencies.

## Discussion

Auxotrophies are widespread among microorganisms (11,34). The obligate nutritional requirements can have far-reaching consequences for the auxotrophic strains and the entire microbial community in the ecosystem (35). On the one hand, each auxotrophy for a specific essential nutrient (e.g., amino acids) increases the organism’s dependence on the nutritional environment, coupling the organism’s survival and proliferation to the availability of the specific compound (35). On the other hand, if the focal metabolite is available, auxotrophic genotypes might gain a selective advantage over prototrophic genotypes by saving metabolic costs (36). In microbial communities, auxotrophies can affect the interactions between microorganisms and their hosts, where auxotrophs could act as recyclers of metabolites that other community members release as by-products of their metabolism (37). In addition, organisms that are auxotrophic for different metabolites could engage in cooperative cross-feeding interactions (38–40). Despite the ecological relevance of auxotrophies, their role in the human gut microbiome is largely unknown. More specifically, Ashniev et al. 2022 showed that several human gut bacterial isolates are indeed amino acid auxotrophs using genome analysis and a comprehensive literature review of experimentally determined auxotrophies and prototrophies (29). Still, the overall distribution and variation of auxotrophies in the human gut microbiome remain elusive. Here, we systematically analyzed the distribution of amino acid auxotrophies in the human gut microbiome using genome-scale metabolic modeling. Moreover, we statistically assessed the associations of inferred auxotrophy frequencies with overall microbiome diversity, long-term stability, and microbial contribution to the human metabolome.

### Ubiquity of auxotrophies indicates a high prevalence of cross-feeding

Overall, high frequencies of auxotrophies were found in the human gut microbiome. For instance, we found that 54%(median) of organisms in the gut microbial communities of healthy adults are auxotrophic for tryptophan. Interestingly, the most frequent auxotrophies for amino acids in the human gut microbiome are also essential nutrients for the human host (Fig. 4A). While auxotrophies in human gut bacteria were reported before, the sources of amino acids for auxotrophic genotypes remain unknown. There are three potential sources of amino acids of auxotrophic members of the gut microbiome:

First, amino acids might be acquired from dietary proteins (41). However, most diet-derived protein is broken down in the upper gastrointestinal tract, and amino acids are absorbed by the human host, limiting protein and amino acid passage to the colon, where most of the gut microbiome resides (41). While most dietary free amino acids do not reach the colon, some dietary proteins that escape digestion in the small intestine can provide a nutrient source for the auxotrophic colonic microbiome(42). Our predictions are based on genomes from stool samples, which predominantly reflect the microbiome composition in the large intestine. Therefore, we argue that the high frequency of amino acid auxotrophies predicted for the colon microbiome in this study is unlikely to be explained by dietary sources of amino acids alone. Plus, we did not find any statistical associations between the dietary intake of amino acids of 79 adults and the frequency of auxotrophies in the microbiome (Supplementary Fig. S7), which further indicates that auxotrophic genotypes acquire their amino acids from other sources. Another study supports our conclusion, as varying dietary concentrations of essential nutrients did not alter the frequency of auxotrophy in the gut (43).

Second, auxotrophs might obtain their essential amino acids by cross-feeding interactions with prototrophic genotypes. Cross-feeding between strains that are auxotrophic for different amino acids has been demonstrated in synthetic (40) and naturally occurring microbial communities (34). Furthermore, a recent study showed that amino acids synthesized by the colonic microbiome stay in the gut and are not absorbed via the mucosa(42). Cross-feeding as a potential source of amino acids for auxotrophic bacteria requires that prototrophic bacteria in the microbial community secrete the respective amino acids. In fact, amino acid biosynthesis and the release into their growth environment have been reported for several gut bacterial species, including members of the genus Bacteroides(44) and the species *Bifidobacterium longum*(45). Thus, cross-feeding enables the growth of auxotrophic organisms even in environments where the focal nutrient is unavailable. Our results suggest a wide diversity of auxotrophic profiles between coexisting genotypes (Fig. 4D), indicating metabolic complementarity and amino acid cross-feeding in gut microbial communities.

Host-derived metabolites are the third potential source of amino acids for auxotrophic gut microbes. Yet, evidence reported in the scientific literature for gut microbial uptake of host-derived amino acids is scarce (42,46). An interesting case where an auxotrophic gut bacterium covers its demand for the focal amino acid might be *Akkermansia muciniphila*. Our predictions show that this bacterium is auxotrophic for threonine, which is in agreement with previous cultivation experiments (15). A. *muciniphila* is a known degrader of host mucins and resides in the mucus layer. Besides glycans, mucin consists of a core protein scaffold rich in proline, threonine, and serine (47). Thus, the threonine auxotrophy of *A. muciniphila* may indicate that this species also utilizes host-derived threonine.

### Auxotrophies might promote ecological diversity and microbiome stability

A major result of our study is the positive associations between auxotrophies and diversity of the human gut microbiome. Earlier studies that used theoretical approaches suggested that auxotrophies can increase and maintain diversity in microbial communities by creating niches for different organisms to occupy through metabolite cross-feeding (25,37). Thus, we conclude that in communities with more auxotrophic members, more cross-feeding may take place, which could promote diversity. Our results support this theory since we observed a positive association between microbiome diversity and auxotrophic profile differences among coexisting genotypes.

Microbe-microbe interactions via metabolite exchanges may also promote microbiome stability (48). Here, we tested if having more auxotrophies as an indicator for metabolite cross-feeding in the gut microbiome is linked to greater stability in healthy adults over three to four years. Indeed, our findings from two independent cohorts indicate that microbiomes with a higher average frequency of auxotrophies at the beginning of the study period remained more stable throughout the duration of the studies (Fig. 5). The association of auxotrophies with microbiome stability was even more pronounced when considering the dissimilarity of auxotrophy profiles of coexisting genotypes as a proxy for amino acid cross-feeding. This result is in line with a theoretical study by Oña and Kost, which demonstrates that cross-feeding between auxotrophs can facilitate that the community structure returns to equilibrium after ecological perturbance (26). Moreover, Sharma et al. (2019) reported that B-vitamin auxotrophies in the human microbiome are prevalent and suggest that cross-feeding B-vitamins between prototrophic and auxotrophic genotypes contributes to gut bacterial population dynamics. The authors also base their conclusion on experimental results, where gnotobiotic mice were colonized by a human fecal microbial community. In these experiments, varying dietary B vitamin intake in mice did not result in appreciable changes in gut microbial community structure, including the proportion of B vitamin-auxotrophic subpopulations, which further suggests cross-feeding as a source of essential nutrients for auxotrophic bacteria in the gut environment and supports our hypothesis that higher auxotrophy frequencies contribute to microbiome stability (Fig 5AB).

Since a reduction in gut microbiome diversity has been reported for several chronic diseases(49–51), our results and the methodology to predict auxotrophy frequencies may guide the development of novel personalized treatment strategies by targeting ecological interactions between coexisting gut microorganisms. For instance, oral administration of microencapsulated amino acids with delayed content release could be used to specifically promote the growth of beneficial subpopulations of the large intestine microbial community, which are auxotrophic for the focal compound (52).

There is an ongoing debate about how different types of cell interactions (i.e., cooperation and competition) contribute to the stability of multi-species communities (20,26,53–55). We want to emphasize that we do not claim that cooperative interactions are stronger than competitive interactions in stabilizing microbiomes, also because we focused in this study on one type of interaction (amino acid cross-feeding) and not on the prevalence of other kinds of interactions or the exchange of other metabolites. Instead, we argue that our results provide evidence that auxotrophies and potential amino acid cross-feeding contribute to maintaining microbiome composition.

### Auxotrophy associations with the human metabolome

Pathways of amino acid biosynthesis and fermentation by-product biosynthesis share common precursors. Therefore, the loss of biosynthetic genes for amino acids might affect the flux distribution in the metabolic network (36). Fermentation by-products such as the organic acids butyrate, acetate, and propionate have implications for human physiology (1). Hence, we wanted to investigate whether specific amino acid auxotrophies are associated with the profile of fermentation products released by gut bacteria. Comparison of the fermentation by-product profile of auxotrophic and prototrophic bacteria revealed statistically significant associations (Fig. 3A), which may be due to the structure of the metabolic network. For example, BCAA auxotrophic bacteria are more likely to be lactate producers, which might be attributed to the fact that the common precursor of BCAA synthesis and lactate synthesis, pyruvate, is no longer used for BCAA synthesis in BCAA auxotrophic bacteria but can be used for lactate formation. The altered fermentation profile in auxotrophic bacteria may, therefore, indicate the importance of the nutritional requirements of gut bacteria for the microbiome’s contribution to the human metabolome. Indeed, when we tested for associations of the relative abundance of amino acid auxotrophs with compounds of the human metabolome, we found several significant correlations (Fig. 4E). In particular, the frequencies of several auxotrophies were correlated with phenylic and indolic metabolites, namely hippuric acid, p-cresol sulfate, 3-indole acetic acid (IAA), and 3-indole propionic acid (IPA). These compounds were previously reported to be of microbial origin or are derived from gut microbially-produced metabolites (56). For instance, hippuric acid and p-cresol sulfate levels were reported to strongly correlate with the microbiome alpha diversity in a large human cohort study (57). P-cresol is known to be produced by gut bacteria that metabolize tyrosine (58), and we found an association with tyrosine auxotrophic gut bacteria. Moreover, the tryptophan-derived IAA is a known agonist of the epithelial human aryl hydrocarbon receptor, an important regulator of intestinal immunity(59). In summary, our results suggest that the contribution of phenylic and indolic compounds to the human metabolome is linked to metabolic processes performed by amino acid auxotrophic gut bacteria.

### Limitations

The method of our study is subject to certain limitations. In our study, auxotrophies were predicted with reconstructed genome-scale metabolic models. Discrepancies between metabolic modelling-based predictions and results from vitro assessments have been reported and discussed previously (13,28,60). Thus, it is crucial to validate *in silico* prediction with *in vitro* results of auxotrophies. Here, we compared our *in silico* results with *in vitro* results for 36 gut bacterial strains and found a sensitivity of 75% for auxotrophy predictions with *gapseq*- reconstructed genome-scale metabolic models. In addition, we performed auxotrophy prediction for 124 genomes from bacterial strains that are not human gut bacteria but known from cultivation experiments to be prototrophic for all 20 proteinogenic amino acids. This test showed that 99.1% of our prototrophy predictions are in line with the experimental data, suggesting that the high prevalence of predicted auxotrophies among the human gut bacterial genomes is not due to a potential technical bias in the *in silico* approach.

## Conclusion

Our study demonstrates the prevalence and impact of auxotrophs in the human gut microbiome. Auxotrophies are common in the human gut microbiome, and interestingly, amino acids essential to the human host are also commonly essential for large fractions of the gut microbiome. Furthermore, human gut microbiomes with high frequencies of auxotrophies were characterized by higher alpha diversity and were more stable over time. Since gut microbial communities commonly display reduced diversity during chronic diseases, auxotrophy frequencies in the human gut microbiome could indicate a healthy gut microbiome. In addition, our results suggest that metabolite cross-feeding networks in gut bacterial communities may be an important factor for stability and maintaining diversity. From a more technical point of view, previous studies have suggested a cautious interpretation of *in silico*-predicted auxotrophies. Therefore, we validated our *in silico* results with experimentally determined auxotrophies reported in scientific literature. This validation indicated the high predictive performance of our method, which used automatic genome-scale metabolic network reconstruction without the need for manual curation of individual genotypes. Thus, the approach can also be applied to microbial communities other than the human gut microbiome.

## Material and Methods

### Reconstruction of genome-scale metabolic models

Genome-scale metabolic models were reconstructed for bacterial genomes from the Human Reference Gut Microbiome (HRGM) genome collection(27,61). The HRGM collection combines isolate and metagenome-assembled genomes (MAGs) from several data sources to summarize genome sequences obtained from human fecal samples. Metabolic models were reconstructed using *gapseq* version 1.2(62). A detailed description of the genome-scale metabolic model reconstruction workflow can be found in the Supplementary Information and Supplementary Table S6.

### Prediction of amino acid auxotrophies

Amino acid auxotrophies were predicted with flux balance analysis(63), where the objective function was set to the flux through the biomass formation reaction. In detail, each model was tested for its ability to form biomass under two different environmental conditions: First, with the growth medium predicted with *gapseq* (see Supplementary Information), and second, with the same medium but where the amino acid of interest was removed. An organism was defined as auxotrophic for a specific amino acid if the organism was able to form biomass in the original medium but not in the medium without the amino acid of interest. Flux balance analysis was performed in R (v4.1.2), the R package *sybil* v2.2.0 (64), and IBM ILOG CPLEX optimizer as linear programming solver. We validated our auxotrophy predictions for 150 organisms (36 from the human gut, 124 known prototrophs from different environments), for which experimental data for amino acid auxotrophies and prototrophies were available in scientific literature (see Supplementary Information for details).

When assessing the distribution of amino acid auxotrophies in sampled individual microbiomes, it is important to consider the relative abundance of different genotypes. To this end, we combined the estimated relative abundances of reference genomes (see ‘Metagenome data processing’) and predicted auxotrophies in the corresponding genomes to calculate the relative auxotrophy abundance *y_j,k_* of amino acid *k* in sample *j* using the equation:

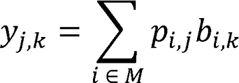

Where *M* is the set of all reference genomes, *p*_*ij*_ the relative abundance of genome; in sample *j*, and *b*_*i,k*_ the auxotrophy prediction with “1” if genotype; is auxotrophic for amino acid *k* and “0” otherwise.

### Prediction of metabolic by-products

For comparison of auxotrophic to prototrophic bacteria, the production rates of fermentation by-product formation were predicted. We undertook this analysis based on the demonstrated accuracy of *gapseq* in predicting fermentation products of anaerobically cultured gut bacteria(62). Given the potential correlation between auxotrophies and the generation of metabolic by-products, investigating auxotrophy distributions could offer new insights into gut microbial metabolism and ecology. Metabolic by-products were predicted with flux-balance-analysis(63) using the flux through the biomass reaction as objective function (i.e., maximization) and subsequently analyzing the fluxes through exchange reactions. Metabolite production rates (mmol*gDW^-1^*hr^-1^) were normalized by growth rates (hr^-1^), resulting in the unit mmol/gDW. Production rates > 1 mmol/gDW were considered as microbial production. The production of the two enantiomers, D-and L-lactate, were combined since their production rates were interchangeable in the FBA solution.

### Cohorts

Data from three human population cohorts were analyzed for the present study. The first cohort comprised paired stool metagenomes and serum metabolomes from 185 participants. This cohort was recruited at the University Hospital Schleswig Holstein, Campus Kiel 2016, and included detailed phenotypic and health-related data. The study was approved by the local ethics committee in Kiel (D441). None of the participants had received antibiotics or other medication two months before inclusion.

The second cohort (Troci et al., 2022) comprised longitudinal stool metagenomes from 79 study participants. Data from this cohort were already part of a previous study (32), which were reanalyzed in the present study. For each participant from this cohort, two metagenomes were sequenced from stool samples that were three years apart. In addition, for each sampling time point, data from food frequency questionnaires were available. In brief, the questionnaire, originally designed and validated for use in the German EPIC study (65), comprised 112 food items and aimed to collect the intake frequency and amount of various types of foods. The average energy intake and other nutrients per day were calculated with data from the German Food Code and Nutrient Data Base (BLS version II.3 (66)). Further information about the sampling method, study design, and sequencing method of the Troci et al. 2022 study can be found in the original publication (32).

The third cohort integrates fecal metagenomes from the 2021 publication by Chen et al., involving 338 Dutch study participants(33). Like the second cohort, the Chen et al. cohort is designed longitudinally, incorporating two fecal metagenomic samples per participant over a four-year interval.

### Metagenome sequencing

DNA of stool samples was extracted using the QIAamp DNA fast stool mini kit automated on the QIAcube (Qiagen, Hilden, Germany) with a prior bead-beating step as described earlier (66). DNA extracts were used for metagenomic library preparation as described previously (32) using Illumina Nextera DNA Library Preparation Kit (Illumina, San Diego, CA) and sequenced with 2×150 bp paired-end reads on a NovaSeq platform (Illumina).

### Metagenome data processing

Metagenomic reads were quality filtered using the ‘qc’ workflow from the metagenome-atlas pipeline tool v2.9.0(67) with default parametrization if not stated otherwise in the Supplementary Information. Quality-controlled (QC) reads were used to estimate the relative abundance of genomes from the HRGM catalog(27) using coverM v0.6.1(68). Across all three analyzed metagenome data sets, a median of 76% QC reads mapped to HRGM reference genomes (Supplementary Fig. S8).

### Targeted metabolomics of blood samples

Metabolite quantification for serum was performed by liquid chromatography tandem mass spectrometry (LC-MS-MS) using the MxP Quant 500 kit (Biocrates Life Sciences AG, Innsbruck, Austria) according to the manufacturer’s instructions. Please refer to the

Supplementary Information document for blood sample preparation and metabolite quantification details.

### Statistical data analysis

All data analysis steps and statistical tests were performed using R (v4.1.2). Flow charts (Fig. 1. and 3A) were created and rendered using Flowchart Designer 3. P-values were corrected for multiple testing using the Benjamini and Hochberg method (69). In all statistical tests, an adjusted p-value of <0.05 was considered as significance threshold. UniFrac distances(70) were calculated using relative abundances of genomes using the R-package abdiv, v0.2.0 (71).

Alpha diversity was calculated using the Shannon index as implemented in the R-package ‘vegan’ v2.6-2 (72). The average pairwise Hamming distance between auxotrophic profiles of co-occurring genomes was calculated per sample to study the effect of metabolic dissimilarity on diversity. In other words, the Hamming distance is the number of amino acids for which the two genotypes had different auxotrophy predictions. In addition to the Hamming distance, we also calculated the abundance-weighted average of auxotrophies per genome y_j_ for each sample j using the equation:

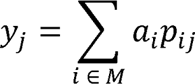

Where *M* is the set of all genomes, *a_i_* the number of auxotrophies in genome;, and *i_ij_* the relative abundance of genome *i* in sample *j*.

For the longitudinal cohorts, the UniFrac distance was correlated with the abundance-weighted average of auxotrophies per genome at the first time point using the Spearman correlation. Further, the Spearman correlation was used to determine the association between the UniFrac distance and the Hamming distance. With food frequency questionnaires, the total dietary intake of amino acids per day was summed up for every individual, and the energy percentage was then calculated based on the total energy intake per day. The Spearman correlation was used to study an association between the total dietary intake of amino acids relative to the total consumed energy (E%) and the frequency of amino acid auxotrophic bacteria. The correlation between the intake of amino acids and frequencies of amino acid auxotrophic bacteria was studied separately for both time points.

## Data availability

The reconstructed genome-scale metabolic models from the HRGM catalog are available via Zenodo (73). Further, metabolic model reconstructions for 124 prototrophic genotypes and 36 gut bacterial genotypes with amino acid auxotrophy/prototrophy status known from laboratory experiments are available via *Zenodo* (74,75). Metagenome sequencing data are provided via the European Nucleotide Archive ‘ENA’ for our study cohort and the cohort from Troci et al. (this study accession: PRJEB60573, Troci et al.: PRJEB48605). Metagenome sequencing data from Chen et al. 2021 (33) are available upon request via the European Genome-Phenome Archive (accession: EGAD00001006959).

## Code availability

The code for analysis of the data can be found in the GitHub repositories https://github.com/SvBusche/Auxo_manuscript_2023 (main results) and https://github.com/Waschina/AGORA2_auxotrophies (for auxotrophy predictions from AGORA2 metabolic models).

## Competing interests

The authors declare no conflicts of interest related to this work.

## Funding

This work was supported by the DFG Cluster of Excellence “Precision medicine in chronic inflammation (PMI)” RTF V and TI-1 (K.A, S.W., P.R.), the PMI Miniproposal funding (S.W.), the EKFS (Clinician Scientist Professorship, K.A.), the BMBF (eMED Juniorverbund “Try-IBD” 01ZX1915A, K.A.), the DFG RU5042 (A.F., K.A., P.R.), DZHK (N.F.), and DFG FR 1289/17-1 (N.F.). This research was further supported through high-performance computing resources available at the Kiel University Computing Centre, which received funding from the German Research Foundation (DFG project number 440395346).

## Acknowledgments

We want to thank the staff of the IKMB (Institute für Klinische Molekularbiologie) microbiome laboratory and the IKMB sequencing facilities for their excellent technical support.

## Author information

### Authors and Affiliations

Institute of Human Nutrition and Food Science, Kiel University, Department of Nutriinformatics, Kiel, Germany

Svenja Starke, Danielle Harris, Silvio Waschina

Institute of Clinical Molecular Biology, Kiel University, Kiel, Germany

Danielle Harris, Mhmd Oumari, Corinna Bang, Philip Rosenstiel, Stefan Schreiber, Andre Franke, Konrad Aden

Research Group Evolutionary Ecology and Genetics, Zoological Institute, University of Kiel, Kiel, Germany

Johannes Zimmermann

Max Planck Institute for Evolutionary Biology, Plön, Germany Johannes Zimmermann

Fraunhofer Institute for Toxicology and Experimental Medicine (ITEM), Hanover, Germany Sven Schuchardt

Department of Internal Medicine III, University Medical Center Schleswig-Holstein, Kiel, Germany

Derk Frank, Norbert Frey

German Centre for Cardiovascular Diseases (DZHK), Partner site Hamburg, Kiel, Lübeck, Germany

Derk Frank

Department of Internal Medicine I, University Medical Center Schleswig-Holstein, Kiel, Germany

Stefan Schreiber, Konrad Aden

### Author contribution

Conceptualization: S.W.

Methodology: Sv.St., Sv.S., S.W.

Software: Sv.St., D.H., S.W., J.Z.

Validation: D.H., S.W., J.Z.

Formal analysis: Sv.St., D.H, M.O.

Investigation: Sv.St.

Resources: Sv.S., D.F., N.F, A.F., K.A., S.W.

Data curation: C.B., M.O., D.F.

Writing - Original Draft: Sv.St., S.W.

Writing - Review & Editing: Sv.St., D.H., S.W.

Visualization: Sv.St., S.W.

Project administration: K.A., S.W.

Funding acquisition: A.F., N.F., P.R., St.S., K.A., S.W.

---

**Table 1.** Cohort characteristics. Age and BMI are given as the median and interquartile.

|  | This study | Troci *et al.*, 2022 | Chen *et al.*, 2021 |
| --- | --- | --- | --- |
| Age (years) | 47 [40-52] | 53 [45.75-57.25]* | 47.5 [40-56] |
| BMI | 24.5 [22.2-26.4] | 25.7 [23.5 -27.5]* | – |
| Female (%) | 44.9 | 37.5* | 55.6 |
| Study participants | 185 | 79 | 338 |

\* Missing values/information: 7.

## Notes

### Competing Interest Statement

The authors have declared no competing interest.

### Summary of Updates

- We included the analysis of an additional and independent cohort study to validate our prior findings concerning the relationship between auxotrophies and microbiome stability. - Extension of the validation of our computational auxotrophy prediction approach. - We reworded and extended large parts of the manuscript to improve clarity in our computational workflow and data analysis description.

