## Supplementary Figure for "Amino acid auxotrophies in human gut bacteria are linked to higher microbiome diversity and long-term stability"

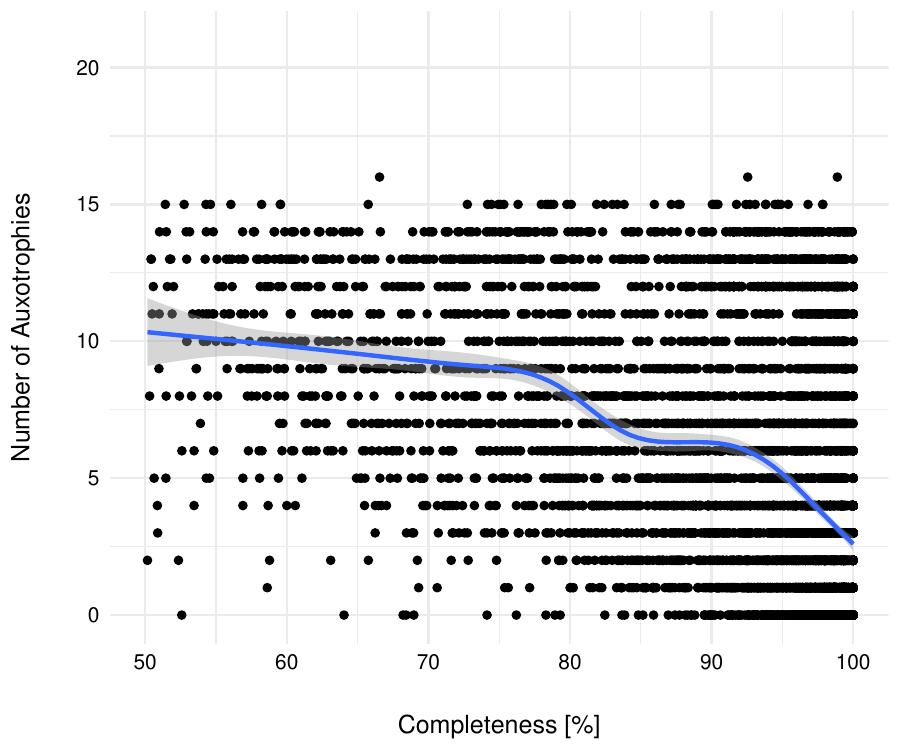

**Supp. Figure S1**: Estimated genome completeness and the predicted number of auxotrophies for 3 687 genomes of representative species from the Human Reference Gut Microbiome (HRGM) collection^1^. The blue line shows the regression line (ρ = -0.50, p ≤ 2.2e-16).

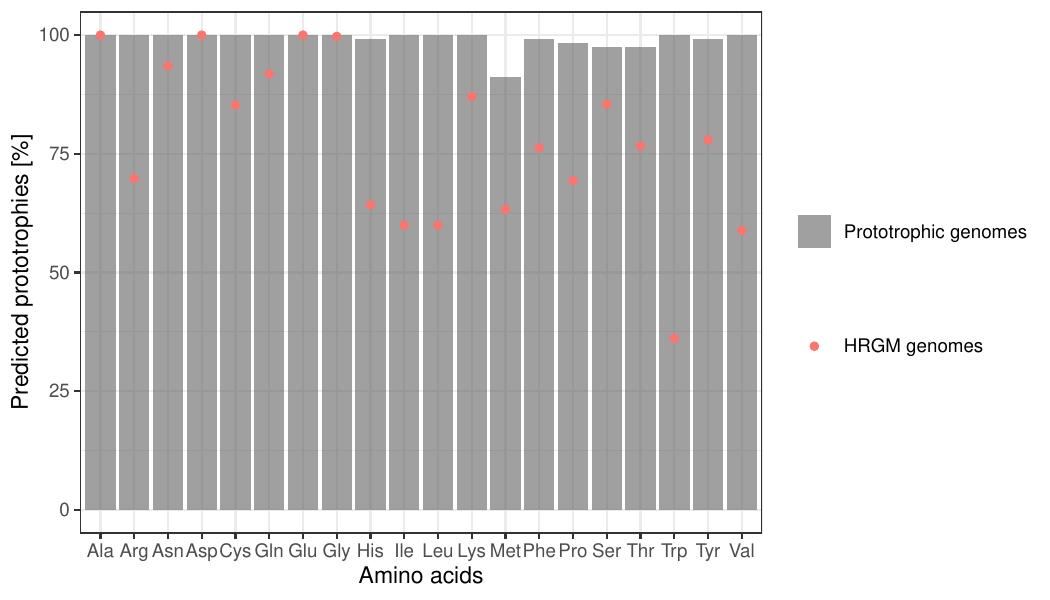

**Supp. Figure S2**: Percentage of *in silico* predicted prototrophies with metabolic modeling in 124 genomes known to be prototrophic^2^ from laboratory experiments (grey bars). The red dots indicate the frequency of prototrophies among 3 687 genomes from human gut bacteria^1^.

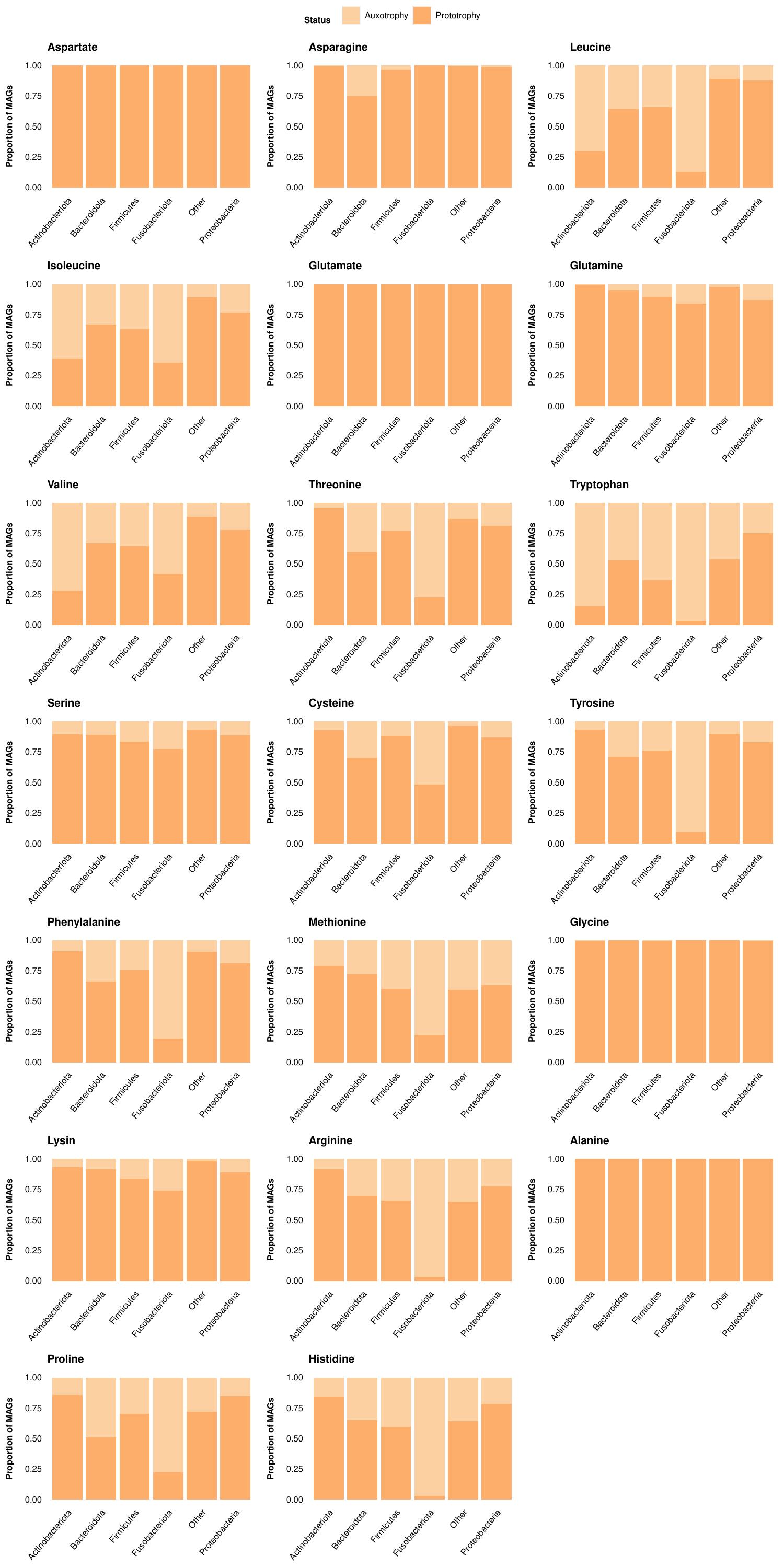

**Supp. Figure S3**: Overview of the proportions of auxotrophy to prototrophy genomes per phylum from the HRGM catalog^1^.

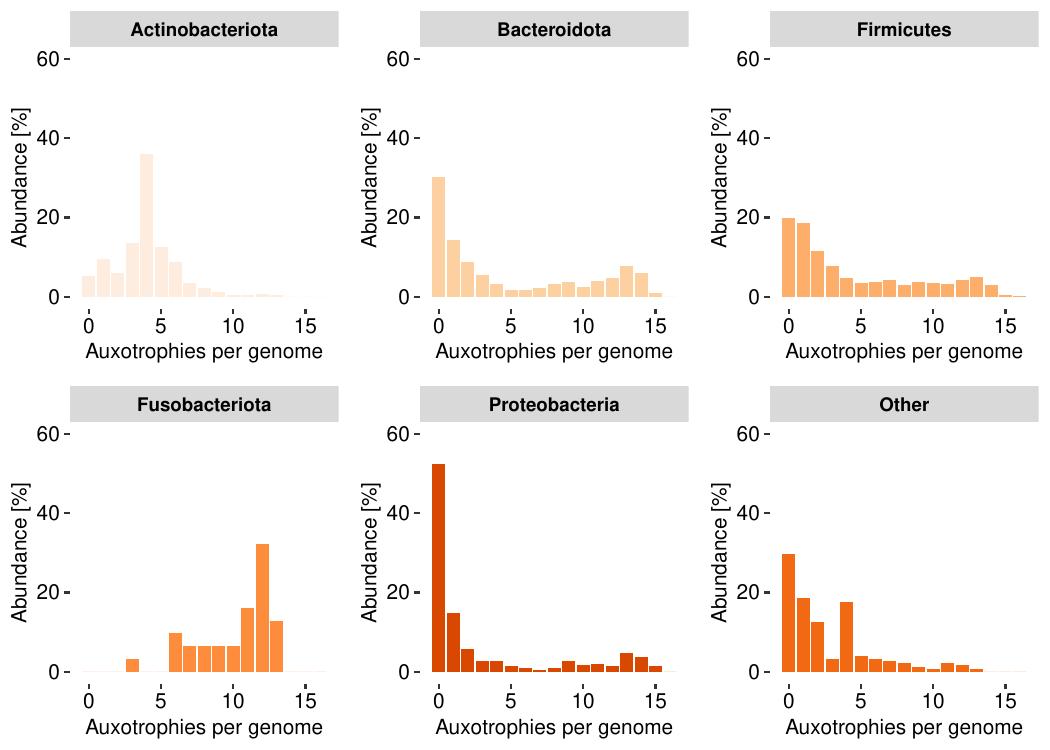

**Supp. Figure S4**: Number of auxotrophies for every phylum. Other is a category that combines different phyla with a lower abundance in the overall HRGM catalogue^1^ and for a reduction of complexity.

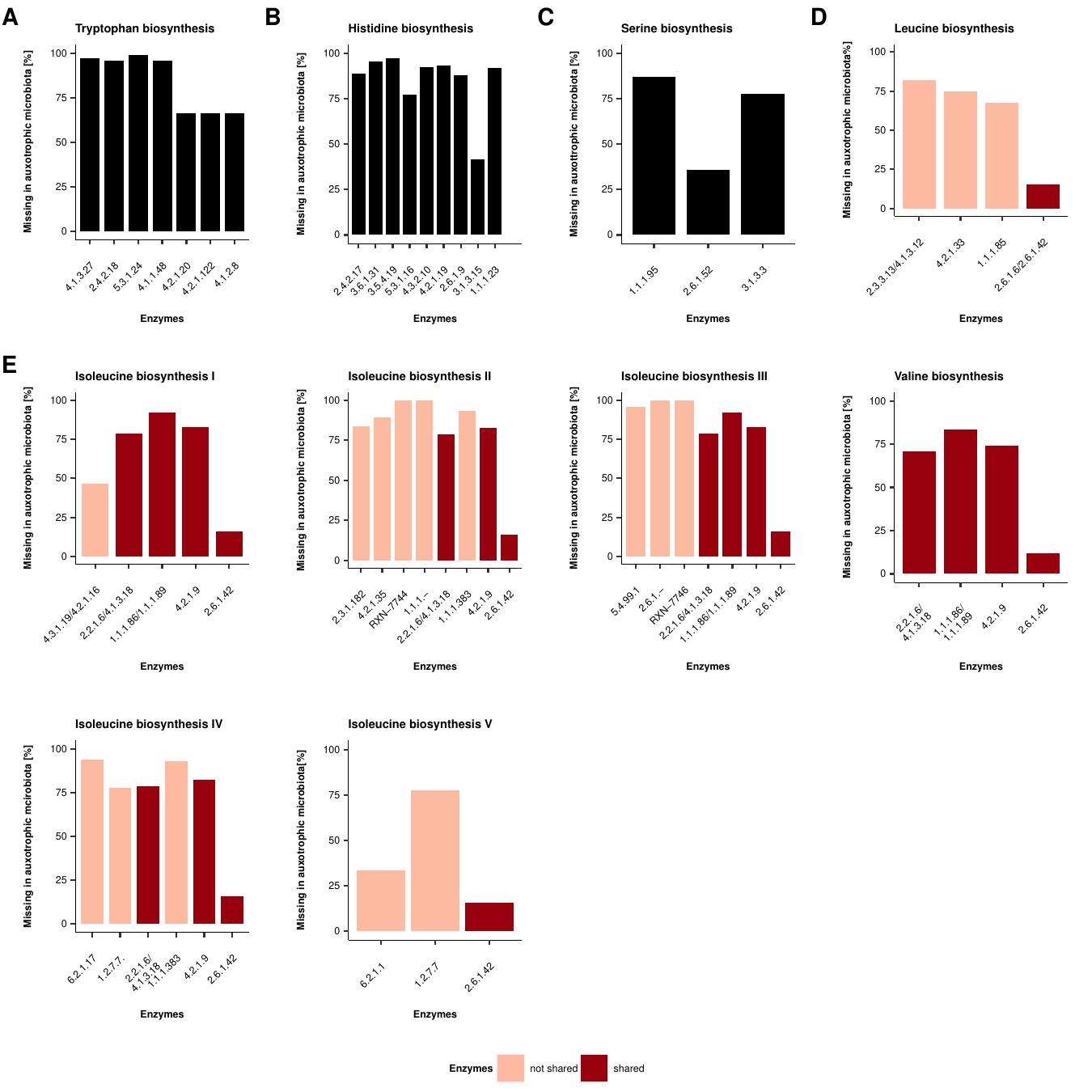

**Supp. Figure S5**: The bar plots display the abundance of missing enzymes in the biosynthesis pathways for several amino acids and BCAA biosynthesis pathways in auxotrophic bacteria. The order of the enzymes in the bar plots represents the one in the pathway. (A) Missing enzymes in the tryptophan pathway of tryptophan auxotrophic bacteria, (B) Missing enzymes in the histidine pathway of histidine auxotrophic bacteria, (C) Missing enzymes in the chorismate pathway of chorismate auxotrophic bacteria, (D) Missing enzymes in the serine pathway of serine auxotrophic bacteria, (E) Comparison of the BCAA pathways of isoleucine, leucine, and valine auxotrophic bacteria, the colors indicate which enzymes are shared in the biosynthesis pathways, the definition of the pathways are based on MetaCyc.

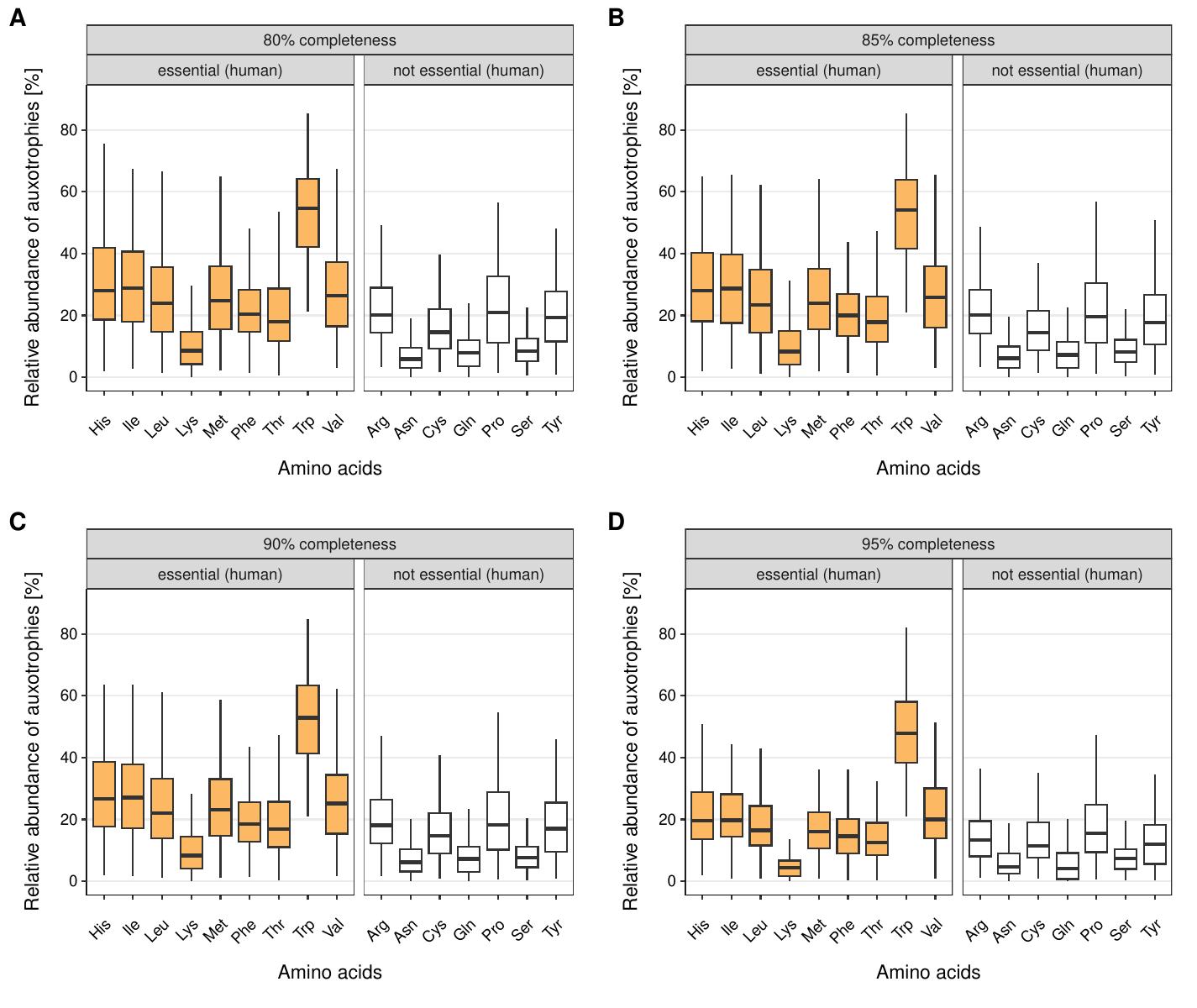

**Supp. Figure S6**: Relative abundance of predicted amino acids auxotrophs depending on the completeness cutoff for reference genomes from the HRGM catalog. Four different genome completeness cutoffs were tested: 80% (A), 85% (B, same as Figure 4A), 90% (C), and 95% (D). The results indicate that the distribution of the relative abundance of predicted amino acid auxotrophies was stable with respect to the chosen completeness cut-off for reference genome filtering.

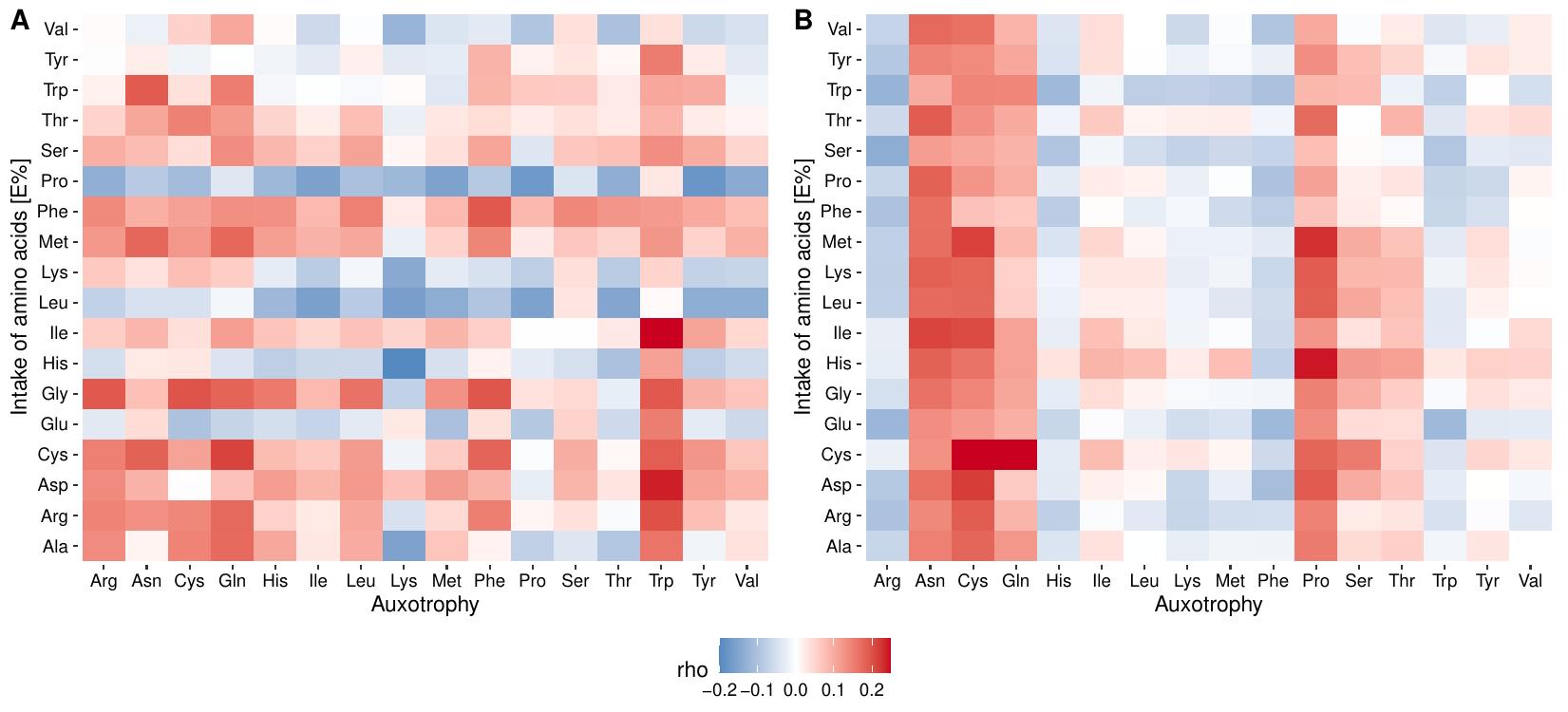

**Supp. Figure S7**: Spearman correlation between the dietary intake of amino acids and the frequency of amino acid auxotrophic bacteria in the gut microbiomes, (A) at the beginning of the study, (B) at the end of the study (3 years later). No statistically significant associations were found (FDR corrected p-value >0.05).

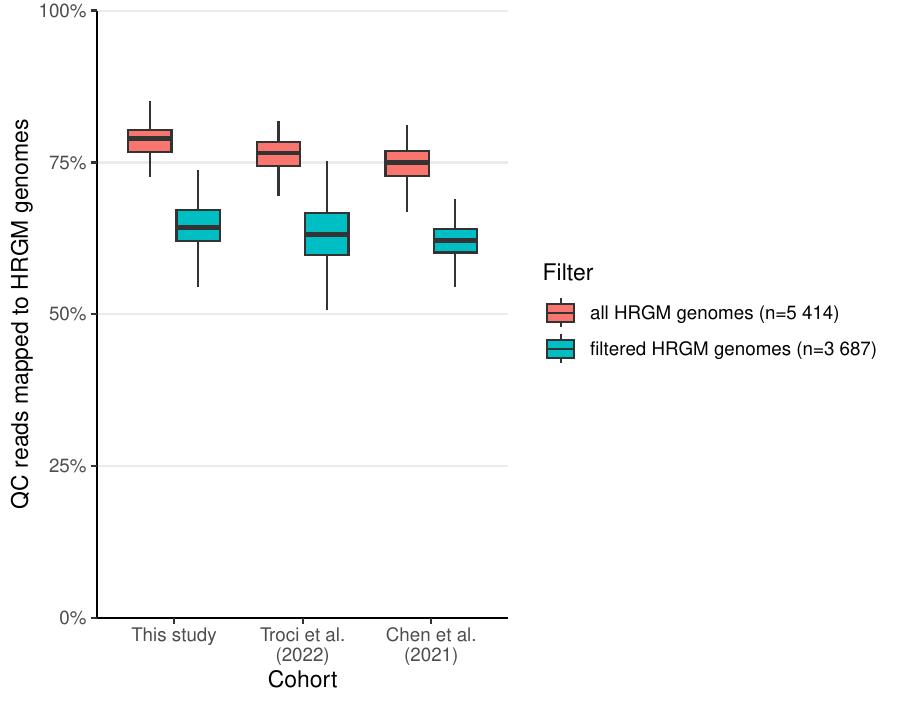

**Supp. Figure S8**: Percentage of quality-controlled metagenomic reads from three cohorts (this study, Troci et al. 2022 ^3^, and Chen et al. 2021 ^4^) to reference genomes from the HRGM catalog.
