## Supplementary Information for "Amino acid auxotrophies in human gut bacteria are linked to higher microbiome diversity and long-term stability"

### Supplementary Material and Methods

**Reconstruction of genome-scale metabolic models**

Genome-scale metabolic models of prokaryotic genomes were reconstructed using *gapseq*^1^. In brief, the *gapseq* reconstruction workflow consisted of five steps: (i) Reaction and pathway prediction, (ii) prediction of metabolite cross-membrane transporters, (iii) reconstruction of a draft metabolic network based on the results from *i* and *ii*, (iv) estimation of an organism-specific growth medium-based on the predicted metabolic capabilities, and (v) gap filling of the metabolic network to enable biomass production using flux balance analysis. Model reconstructions were limited to bacterial genomes marked as representative species in the HRGM collection. Further, genomes with an estimated contamination percentage of ≤2% or a completion ≥85% were included. Based on these filters, 3 687 bacterial genomes were subject to metabolic model reconstruction. Among those genomes, 22% are from bacterial isolates, and 78% are metagenome-assembled genomes.

Borer and Magnúsdóttir have recently shown in a systematic analysis of isolate- and metagenome-assembled genomes that the gap-filling medium strongly impacts the auxotrophies predicted by genome-scale metabolic modelling^2^. We note that with the reconstruction procedure used in this study, we do not rely on an arbitrarily defined gap-filling medium, which is used for every metabolic network model. Instead, for each draft network, a genome-specific gap-filling medium is predicted (see section “Prediction of a genome-specific gap-filling medium” below for details). In brief, if the medium prediction algorithm of *gapseq* finds a known biosynthetic pathway for a specific amino acid in the draft metabolic network, the amino acid will not be part of the resulting predicted medium, as the compound is likely not required from the growth environment, thus, also not necessary for subsequent gap filling. In contrast, if the medium prediction algorithm does not detect at least one complete known biosynthetic pathway for a specific amino acid, the respective compound is added to the outcome gap-filling medium based on the rationale that this amino is a putative essential compound that needs to be obtained from the growth environment. However, it is important to note that this case does not directly imply that the organism is auxotrophic for the specific amino acid, as the subsequent gap-filling algorithm might add reactions to the model that complete a biosynthesis route from other compounds in the gap-filling medium to the amino acid. Such reactions are only added in cases where a gene was found in the query genome that displays sequence similarity to a reference gene sequence with the respective enzymatic function but where the sequence similarity was not high enough to pass the threshold of the bitscore 200 to be directly included in the draft network. For details on the gap-filling algorithm implemented in gapseq, please refer to the original gapseq publication^1^.

Taken together, the model reconstruction and auxotrophy prediction that we used for the present study do not depend on one gap-filling medium composition that is defined for all organisms but adjusts the medium for each organism based on its genome information and by using the multi-step gap-filling algorithm that is implemented in *gapseq*.

**Prediction of a genome-specific gap-filling medium**

The genome-scale metabolic network reconstruction process of *gapseq* requires a gap-filling medium for the final gap-filling step (see above). Here, we used a medium prediction feature (module “*gapseq medium*”) of the gapseq software, which can be plugged into the reconstruction workflow between the homology-based generation of a draft metabolic network and the gap-filling algorithm. We provided the additional command line option “-c cpd00007:0” to ensure that the predicted medium does not contain oxygen (compound identifier: cpd00007). The algorithm for medium prediction tests which pathways and reactions are absent or present in the draft metabolic model. Whether a particular compound is added to the medium is decided using logical expressions that include variables for the presence (TRUE) and absence (FALSE) of pathways and reactions within the design network (see Supplementary Table S6 for all compounds and their logical expressions). For example, the disaccharide lactose (ModelSEED ID: cpd00208), has the logical expression ("LACTOSECAT-PWY" | "LACTOSEUTIL-PWY" | "BGALACT-PWY"), which means, that lactose is added to the medium if one of three known lactose degradation pathways as defined in MetaCyc^3^ is already present in the draft network. In particular, for amino acids, the medium prediction module uses a similar approach to the auxotrophy prediction tool GapMind^4^, which tests if a known biosynthetic pathway for a specific amino acids exists based on sequence homology. The amino acid biosynthetic pathways that are considered are also those that are defined in MetaCyc^3^. For instance, if none of the five known biosynthesis pathways in prokaryotes for lysine (https://metacyc.org/META/NEW-IMAGE?type=PATHWAY&object=LYSINE-SYN) is found, lysine is added to the gap-filling medium.

The medium prediction module of gapseq considered 74 compounds (Supplementary Table S6), including inorganic compounds, carbohydrates, amino acids, other carboxylic acids, and vitamins. Most of those potential nutrients are also compounds that can be found in the growth environment of colonic microorganisms, such as fibers (e.g., pectin, inulin), other dietary compounds (e.g., sulfoquinovose, daidzein), constituents of the mucins (e.g., *N*-acetylneuraminate, *N*-acetylneuraminate) and inorganics (e.g., H_2_, H_2_S, H_2_O).

Besides the enumeration of available nutrients, a gap-filling medium requires their individual maximum uptake rates by the microorganism. The medium prediction implemented in *gapseq* uses rates commonly used in manually curated genome-scale metabolic network models, e.g., 0.1 mmol*gDW^-1^*hr^-1^ for amino acids or 5 mmol*gDW^-1^*hr^-1^ for monosaccharides. In the case of oligo- and polysaccharides, the maximum uptake rates are scaled to allow the same uptake rate per subunit (e.g., 5 mmol*gDW^-1^*hr^-1^ for the monosaccharide D-glucose and 2.5 mmol*gDW^-1^*hr^-1^ for the disaccharide maltose).

**Validation of auxotrophy predictions**

To validate our *in silico* predicted auxotrophies, we collected the genome sequences from NCBI RefSeq for 36 bacterial strains for which experimental data were available on amino acid auxotrophies/prototrophies. The majority of these strains were already summarized by Ashniev et al. 2022 ^5^. In that study, the authors even list more than the 36 strains we analyzed here. This is because we excluded cases where we could not find a genome assembly of the exact strain used in the referenced experimental study. Moreover, we excluded the entries for species belonging to the genus *Bifidobacterium* since their auxotrophy for cysteine is ambiguous: Some studies report cysteine as an essential nutrient for growth^6,7^, while genomic analysis and genome-scale metabolic modeling indicated the presence of cysteine biosynthetic pathways^8,9^. This ambiguity most likely stems from the fact that cysteine biosynthesis in *Bifidobacterium* species depends on the available sulfur source^8^. Genomic analysis of *Bifidobacterium bifidum* PRL2010, for instance, suggested the strain’s inability to use sulfate as a sulfur source, while hydrogen sulfide or methionine could potentially serve as a sulfur source for the biosynthesis of cysteine^6,8^.

Genome-scale metabolic models for all 36 strains were reconstructed as described above. Auxotrophies were predicted with the method described in the main manuscript.

In addition to the gapseq-reconstructed models, we also predicted auxotrophies for 20 of the 36 bacterial strains using genome-scale models from the AGORA2 collection^10^ (Supplementary Table S2). Auxotrophies were predicted in the same manner as for gapseq models. In contrast to the gapseq models, which can contain only free amino acids and not peptides in the predicted medium, some AGORA2 models have exchange reactions with lower bounds < 0 for dipeptides. In those cases, and for predicting the auxotrophy status for amino acid x, we changed the lower bound to 0 for all exchange reactions of dipeptides involving amino acid x. At the same time, we introduced a new inflow reaction of the non-x amino acid moiety to the model to predict only the essentiality of x and not of the other amino acids in the respective peptides. The sensitivity, specificity, and accuracy of auxotrophy predictions were calculated for *gapseq* models (n=36) and AGORA2 models (n=20).

As an additional auxotrophy prediction validation step, 124 genome-scale metabolic models were reconstructed for bacterial strains that were reported by Price, 2023, to be able to grow in a defined growth medium containing no amino acids^11^. Thus, these 124 organisms are known amino acid prototrophs and can be used to estimate the rate of false auxotrophy predictions. The original publication by Price reported 127 genomes of prototrophs; however, 3 of the corresponding genome assemblies (GCF_000014265.1, GCF_000020545.1, GCF_900188395.1) were suppressed on RefSeq at the time we performed the analysis in March 2023. Auxotrophies for the 124 genome assemblies of this prototroph collection were predicted as described above, and results are summarized in Supplementary Table S3.

**Metagenome data processing**

Metagenomic reads were subject to quality control and filtering using the ‘qc’ workflow from the metagenome-atlas pipeline tool v2.9.0^12^. In detail, reads were (1) deduplicated, (2) quality filtered, and (3) decontaminated. Modules from the BBmap suite v37.99 (BBMap - Bushnell B. - sourceforge.net/projects/bbmap/) were used for all three steps. In the deduplication step (1), the BBmap module *clumpify.sh* was used with the parameters “dedupe=t dupesubs=2”, which removed duplicate reads with a maximum of 2 substitutions between duplicates. The quality filter (2) employed the BBmap module *bbduk.sh* with the parameters “hdist=1 ktrim=r mink=8 trimq=10 qtrim=rl minlength=51 maxns=-1 minbasefrequency=0.05” and otherwise default options. This quality filter trimmed reads from the right if adapter sequences were detected, trimmed reads on both sides from the first base with a quality score below 10, removed sequences that were shorter than 51 bp after trimming, removed sequences with ambiguous base calls (i.e., “N”s), and removed reads if any base had a frequency of less than 5%. Finally, reads that are likely contaminations from the human host genome or Illumina PhiX sequences were removed using the BBmap module *bbsplit.sh* using the option “maxratio=0.65” and otherwise default parametrization. This tool tested if specific reads mapped to the host genome or PhiX sequences based on sequence similarity and mapped reads were discarded from the sample’s fastq files. For the decontamination step, the human reference genome assembly ‘Genome Reference Consortium Human Build 38’ (GRCh38) was used, in which low entropy regions (entropy < 0.7) were masked using the *bbmask.sh* tool within the BBmap suite. Moreover, regions that display high similarity to prokaryotic rRNA genes were additionally masked. To this end, prokaryotic small and large subunit rRNA gene sequences were retrieved from SILVA version 138.1^13^ and shredded into shorter (80 bp) sequences with 40 bp overlaps using *shred.sh*. Shredded sequences were aligned to GRCh38 with a minimum identity of 85% and maximum indel length of 2 bp. Regions in GRCh38 with alignment hits were masked.

As mentioned in the main manuscript, we used the Human Reference Gut Microbiome ‘HRGM’ catalog^14^ as reference genomes for quantifying representative microbial genomes in the metagenomic data sets. We had chosen this collection, as it was the latest published human gut microorganism genome collection when we were finalizing the results of the present study. Furthermore, the HRGM collection contains 780 species-level representative genomes, which were absent in previous genome collections and assembled from metagenome samples from before under-represented Asian countries, namely Korea, Japan, and India. For each metagenome sample, the relative abundance of HRGM genomes was estimated using coverM^15^ v0.6.1 with default parametrization of the module `coverm genome`.

**Targeted metabolomics of blood samples**

Serum samples were collected using serum s-monovette (9ml, Sarstedt, Germany). Samples were incubated upright at RT for 30 min. and centrifuged (10 min., 2000 x g). Serum was aliquoted in 500μl tubes and stored at -80°C. Metabolite quantification for serum was performed by liquid chromatography tandem mass spectrometry (LC-MS-MS) using the MxP Quant 500 kit (Biocrates Life Sciences AG, Innsbruck, Austria) according to the manufacturer's instructions. The MxP Quant 500 kit simultaneously measures 630 metabolites covering 14 small molecule and 12 different lipid classes. It combines flow injection analysis tandem mass spectrometry (FIA-MS/MS) using SCIEX 5500 QTrap mass spectrometer (SCIEX, Darmstadt, Germany) for lipids and liquid chromatography tandem mass spectrometry (LC-MS/MS) using Agilent 1290 Infinity II liquid chromatography (Santa Clara, CA, USA) coupled with a SCIEX 5500 QTrap mass spectrometer for small molecules using multiple reaction monitoring (MRM) to detect the analytes. Data evaluation for serum metabolite concentrations and quality assessment was performed with the software SCIEX Analyst software (Version 1.7.2) and the MetIDQ™ software package (Oxygen-DB110-3023), which is an integral part of the MxP Quant 500 kit.

For downstream statistical analysis, the serum metabolome data were pre-processed by imputing missing specific values using a random forests approach as implemented in the R-package ‘missForest’ and the function with the same name in default parametrization ^16^. This imputation was limited to missing values for metabolites, which have less than 20% missing values across the data set.

With a partial Spearman correlation, the association between the frequency of auxotrophic bacteria and serum metabolites and other hematology parameters from the DZHK cohort was evaluated ^17^. We adjusted for the potential confounders sex, age, and BMI. P-values were corrected for multiple testing using the False Discovery Rate (FDR) method.
